# Localized K63 ubiquitin signaling is regulated by VCP/p97 during oxidative stress

**DOI:** 10.1101/2024.06.20.598218

**Authors:** Austin O. Maduka, Sandhya Manohar, Matthew W. Foster, Gustavo M. Silva

## Abstract

Under stress conditions, cells reprogram their molecular machineries to mitigate damage and promote survival. Ubiquitin signaling is globally increased during oxidative stress, controlling protein fate and supporting stress defenses at several subcellular compartments. However, the rules driving subcellular ubiquitin localization to promote these concerted response mechanisms remain understudied. Here, we show that K63-linked ubiquitin chains, known to promote proteasome-independent pathways, accumulate primarily in non-cytosolic compartments during oxidative stress induced by sodium arsenite in mammalian cells. Our subcellular ubiquitin proteomic analyses of non-cytosolic compartments expanded 10-fold the pool of proteins known to be ubiquitinated during arsenite stress (2,046) and revealed their involvement in pathways related to immune signaling and translation control. Moreover, subcellular proteome analyses revealed proteins that are recruited to non-cytosolic compartments under stress, including a significant enrichment of helper ubiquitin-binding adaptors of the ATPase VCP that processes ubiquitinated substrates for downstream signaling. We further show that VCP recruitment to non-cytosolic compartments under arsenite stress occurs in a ubiquitin-dependent manner mediated by its adaptor NPLOC4. Additionally, we show that VCP and NPLOC4 activities are critical to sustain low levels of non-cytosolic K63-linked ubiquitin chains, supporting a cyclical model of ubiquitin conjugation and removal that is disrupted by cellular exposure to reactive oxygen species. This work deepens our understanding of the role of localized ubiquitin and VCP signaling in the basic mechanisms of stress response and highlights new pathways and molecular players that are essential to reshape the composition and function of the human subcellular proteome under dynamic environments.

## INTRODUCTION

Oxidative stress is involved in the progression of many diseases, including cancers, infections, cardiovascular diseases, and neurodegenerative diseases (1). Oxidative stress arises from the overaccumulation of reactive oxygen species (ROS), leading to molecular damage and often causing apoptosis or necrosis (2). In response to elevated ROS, human cells promote survival through rewiring regulatory events for transcription, translation, protein degradation, and overall signal transduction (3–5). Several of these events are controlled by dynamic and reversible post-translational protein modifications (PTMs) that are necessary for survival in response to oxidative stress (6, 7). In this context, PTMs serve in various roles to promote cellular defense, such as regulating degradation of damaged proteins by the proteasome, or reprogramming transcription and translation to activate stress-responsive genes and produce antioxidant proteins (8–12). Although genomic, proteomic and antibody technologies have improved our knowledge on the complexity of PTMs (13), many roles of post-translational modifications in the context of oxidative stress remain unexplored.

Ubiquitination is a diverse and expansive PTM of proteins in eukaryotes, with over 70,000 ubiquitinated peptides identified from over 8,000 proteins regulating virtually every cellular pathway (14). Studies on ubiquitination have initially focused on its role in proteasomal degradation, however ubiquitin can form distinct chain architectures that provide a broader range of functions (15–17). The ubiquitin monomer has seven lysine residues and a methionine (M1) residue that can by modified by ubiquitin itself to form unique polyubiquitin chains (18). For example, lysine-63 linked polyubiquitin chains (K63ub) have been implicated in the oxidative stress response in yeast, and K27ub has been shown to be essential for mammalian cell proliferation (6, 19). These various linkages are formed by families of ubiquitin conjugating enzymes (E2s) and ubiquitin ligases (E3s) and reversed by deubiquitinases (DUBs) (20). Subsequently, unique ubiquitin linkages on substrates can be recognized by proteins containing ubiquitin binding domains (UBDs) that help process these chains and facilitate downstream signaling (21). Many of these processing factors that trigger distinct signaling pathways have been shown to be differentially regulated during oxidative stress (10, 22). Still, mechanisms involving proteasome-independent ubiquitination during oxidative stress remain understudied.

Many components of the ubiquitin system are regulated at the subcellular level (23–25). For example, K48ub linkages are utilized for protein quality control in the endoplasmic reticulum (ER) via ER-associated degradation (ERAD). Local dysregulated populations of ribosomes are modified by ubiquitin in many stress responses (26), such as K63ub in ER-localized ribosome quality control (RQC) (9). ER-localized K63 ubiquitination of G3BP1 is involved in the disassembly of heat-induced stress granules (27). K27ub is important for normal nuclear dynamics and cell cycle progression (19), and K63ub can also modify plasma membrane proteins to facilitate their endosomal-to-lysosomal sorting (25). Ubiquitin modifications on proteins are generally elevated at the whole-cell level in response to oxidative stress (28), but we are just uncovering how this increase is resolved on a subcellular level, and how localization contributes to response pathways activated under oxidative stress. Localization of specific ubiquitin linkages has been studied with microscopy using fluorescence-based UBD sensors (29), however these approaches are limited in resolving the molecular substrates and factors involved.

In both subcellular compartments and at the whole cell level, proteomic techniques have been widely used to study protein ubiquitination (30–32). Enrichment and isolation methods revolutionized the field and allow researchers to decipher the complex network of ubiquitin signals in dynamic environments (33, 34). These techniques have been employed in human cells to study ubiquitination in a variety of stress environments (35). However, deeper insights can be uncovered utilizing proteomics to study the subcellular localization of ubiquitin signaling in the context of oxidative stress. In this study, we discovered that K63ub signals selectively and reversibly accumulate in non-cytosolic compartments in response to oxidative stress induced by sodium arsenite (NaAsO_2_) in human cells. We also use ubiquitin peptide enrichment proteomics to uncover details of the ubiquitin subcellular landscape as well as downstream effectors under oxidative stress, linking non-cytosolic signaling to immune responses and translation. We found that VCP/p97 and its adaptor NPLOC4 similarly accumulate and are critical for a cyclical processing of proteins linked to K63ub chains. Our work highlights the regulation and complexity of ubiquitin signaling in humans under stress, and provides an approach to comprehensively study localized ubiquitin responses under harmful environments.

## RESULTS

### Arsenite stress induces subcellular partitioning of K63 ubiquitin signals

To understand if ubiquitin signaling is spatially distinct under oxidative stress, we used a detergent-based method to enrich for cytosolic and non-cytosolic components—including membranes and organelles—after NaAsO_2_ treatment (**Fig. 1A**). Arsenite is an oxidative stressor that generates ROS through mitochondrial decoupling (36). Because of its proteasome-independent function (37), we analyzed the dynamics of K63-linked ubiquitin chains, which are amongst the most abundant linkage types in humans (38). We observed in HeLa cells that K63ub, but not K48ub, accumulates primarily in non-cytosolic compartments of cells after 1 hour exposure to 0.5mM NaAsO_2_ (**Fig. 1B**). These findings were reproduced in HEK293T and U2OS cells (**supplemental Fig. S1A-S1B**), suggesting a conserved mechanism of K63ub signal partitioning across cell lines. To further validate that K63ub signals selectively accumulate in non-cytosolic compartments, we transfected cells with HA-tagged ubiquitin constructs and found that a mutant with all lysine residues mutated to arginine except lysine-63 still accumulated (**Fig. 1C**). To further confirm the K63ub accumulation is occurring outside of the cytosol, we used confocal microscopy merged with a similar detergent-based method (39) to remove cytosolic components of our samples before fixation (**Fig. 1D**). First, we found that GAPDH, a cytosolic marker, is removed by digitonin treatment. Moreover, we observed that K63ub puncta is significantly formed upon oxidative stress, which remained present after digitonin treatment but are absent after non-cytosolic compartment removal with DDM (**Fig. 1D-E**). To further confirm accumulation of K63ub is dependent on arsenite, we transferred cells to fresh media to allow them to recover from stress. We found that after eight hours of recovery, K63ub chains return to steady-state levels (**Fig. 1F**), highlighting the reversibility of this signal. Supporting this, formation of non-cytosolic K63ub puncta is also reversible during stress recovery (**Fig. 1G**). As K63ub signals accumulate mainly in non-cytosolic compartments, we investigated whether they accumulate directly in non-cytosolic compartments or whether ubiquitination occurs in the cytosol first prior to being trafficked to other cellular compartments. Therefore, we performed an arsenite time-course experiment to observe cytosolic ubiquitin dynamics during a one-hour NaAsO_2_ treatment. No major accumulation of cytosolic K63ub was seen along the time course compared to non-cytosolic (**Fig. 1H**), suggesting non-cytosolic changes in ubiquitination occurs in a localized fashion. Collectively, our data reports a linkage-specific, conserved, subcellular accumulation of K63ub signals in non-cytosolic compartments in response to arsenite stress.

**Figure 1.**
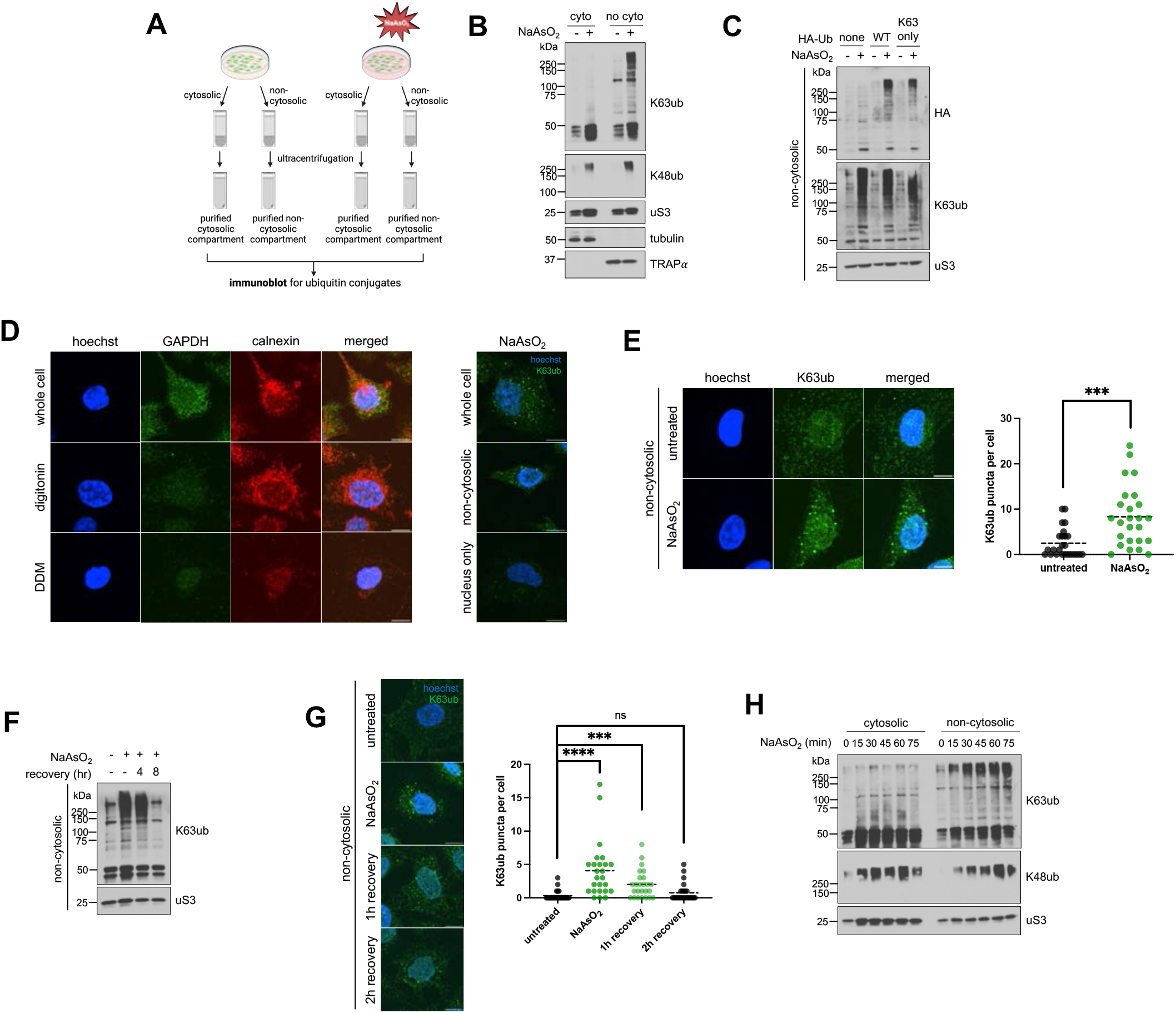
Arsenite stress promotes reversible non-cytosolic K63 ubiquitin accumulation. 1A – Schematic of the cytosolic and non-cytosolic fractionation and sucrose sedimentation approach. Cells were treated as described, then detergents were used to extract cytosolic fractions, followed by non-cytosolic fractions from cells attached to plate. Then fractions were then further purified by ultracentrifugation over a sucrose cushion. 1B – Arsenite stress induces local accumulation specifically for K63 ubiquitin signals in purified non-cytosoilc compartments. Immunoblots of anti-K63ub and anti-K48ub from fractions of cells treated with 1 hour of 0.5 mM NaAsO_2_. Anti-uS3 was used as a loading control for each fraction. Before ultracentrifugation, anti-tubulin and anti-TRAPα were used as fractionation controls for the cytosol and non-cytosolic fractions, respectively. 1C – Exogenous HA-Ub(K63) accumulates in the non-cytosolic fraction upon arsenite stress. Immunoblot of anti-HA and anti-K63ub from non-cytosolic fractions of cells transfected with either WT or K63-only HA-ubiquitin plasmids, then treated with 1 hour of 0.5 mM NaAsO_2_. Anti-uS3 was used as a loading control. 1D - Approach for observing non-cytosolic K63ub puncta dynamics. Immunofluorescence microscopy of anti-GAPDH (cytosol) and anti-calnexin (ER/membrane) from HeLa cells prepared either as whole cells, cytosol-depleted cells after a digitonin wash, or membrane-depleted cells after a digitonin and DDM wash. Hoechst was used to identify and count cells. Protocol was repeated on the right after treatment of 1 hour 0.5mM NaAsO_2_, then stained for anti-K63ub. Hoechst was used to identify cells. Scale bar = 10 *μ*m. 1E – Non-cytosolic K63ub linkages form puncta upon arsenite stress. Immunofluorescence microscopy of anti-K63ub from cells treated with 1 hour of 0.5 mM NaAsO_2_. Cells were prepared with a digitonin wash to release cytosolic components before fixation. Hoechst was used to identify and count cells. Analysis included 25 cells per group. Scale bar = 10 *μ*m. 1F – Non-cytosolic K63ub accumulation is reversible after arsenite washout. Immunoblot of anti-K63ub from non-cytosolic fractions of cells along a course of treatment with 1 hour of 0.5 mM NaAsO_2_ with a washout and recovery of either 4 or 8 hours. Anti-uS3 was used as a loading control. 1G – Non-cytosolic K63ub puncta formation is reversible after arsenite washout. Immunofluorescence microscopy of anti-K63ub from cells along a course of treatment with 1 hour of 0.5 mM NaAsO_2_ with a washout and recovery of either 1, 2, or 4 hours. Cells were prepared with a digitonin wash to release cytosolic components before fixation. Hoechst was used to identify and count cells. Analysis included 25 cells per group. Scale bar = 10 *μ*m. 1H - Arsenite stress does not induce accumulation of K63 ubiquitin signals in purified cytosolic fractions. Immunoblots of anti-K63ub and anti-K48ub from fractions of cells treated along a 15-minute time course of 0.5 mM NaAsO_2_ up to 75 minutes. Anti-uS3 was used as a loading control for each purified fraction. Before ultracentrifugation, anti-tubulin and anti-TRAPα were used as fractionation controls for the cytosol and non-cytosolic fractions, respectively.

K63ub signals have been implicated in many organellar and membrane-associated pathways downstream from stress events like ER stress and ribosome collisions (40). Therefore, we sought to determine whether this subcellular accumulation of K63ub signals is due to these canonical stress responses that are inducible by oxidizing environments. ER stress is an overload of misfolded ER proteins that can trigger degradation pathways to restore proteostasis (41). We found that direct induction of ER stress by tunicamycin incubation (41) does not lead to similar K63ub accumulation as arsenite treatment (**supplemental Fig. S2A**), suggesting ER stress is not responsible for the accumulation of K63ub chains. RQC also involves ribosome K63-linked ubiquitination during stress in response to ribosome collision events, which can be induced by low levels of the translation inhibitor anisomycin (40). We also found that a low dose of anisomycin that induces ribosome collisions does not induce high levels of K63ub accumulation (**supplemental Fig. S2B**). Moreover, K63 ubiquitination of ER-localized G3BP1 is involved in the disassembly of heat-induced stress granules (27). Still, depleting G3BP1 did not prevent K63ub accumulation during arsenite stress (**supplemental Fig. S2C**). G3BP1-based stress granules also did not colocalize with arsenite-induced non-cytosolic K63ub puncta, nor were they primarily in non-cytosolic compartments under stress (**supplemental Fig. S2D-S2E**). These data are consistent with literature that suggests heat and oxidative stress elicit different changes in the ubiquitin landscape (35). Finally, we depleted E2 ubiquitin enzymes known to be involved in K63ub-mediated pathways (10, 42), and we showed that neither knockdown nor inhibition of UBE2A (10) or UBE2N (42) drastically impaired K63ub accumulation during stress (**supplemental Fig. S3A-S3C**). These data support that K63ub partitioning is not solely dependent on global stress response pathways suggesting that additional mechanisms contribute to this signaling pathway in a subcellular distributed fashion.

### Proteomics reveals stress-induced subcellular ubiquitination and recruitment of processing factors

After highlighting that additional or combinations of mechanisms might contribute to spatial regulation of K63ub signals under oxidative stress, we sought to elucidate the nature of proteins involved in this subcellular response. Here, we used quantitative mass spectrometry to identify both ubiquitinated proteins that accumulate in non-cytosolic compartments under stress, and ubiquitin factors that could affect the dynamics and processing of K63ub substrates. To identify ubiquitin substrates, we performed ubiquitin proteomic analysis in cytosolic and non-cytosolic compartments of cells treated with or without NaAsO_2_. After fractionation, samples were trypsin digested, and ubiquitinated lysine residues left a di-glycyl (K-GG) remnant. These K-GG remnants on tryptic peptides were enriched for using an immunoaffinity approach (**Fig. 2A**) and used to quantify ubiquitinated proteins by mass spectrometry with amino acid resolution (43). After enrichment, LC-MS/MS was performed with a Evosep One LC coupled to a Thermo Orbitrap Astral via a Nanospray Flex ionization source, using a data-independent acquisition method. Principal component analyses of these datasets revealed clustering of all replicates by the presence or absence of stress as well as by compartment, adding to our confidence in the data (**supplemental Fig. S4A-S4B**). Amongst all compartments and treatments, we identified 25,567 unique ubiquitinated peptides, corresponding to 3,967 unique proteins (**supplemental Table S1-S2**). When considering their localized presence, we observed that 15,747 peptides of 3,191 ubiquitinated proteins were at least 1.5-fold enriched in the non-cytosolic compartment (**Fig. 2B**). In the presence of arsenite, we determined that 9,421 peptides out of 2,046 proteins (around 35% of the total dataset) were significantly ubiquitinated under stress in the non-cytosolic compartment with a 1.5-fold change, indicating the range of potential targets that are selectively modified by ubiquitin. By comparing with a focused ubiquitin proteomics study using sodium arsenite as a stressor (35), we observed that 74% of their dataset is shared with ours (**Fig. 2C**). However, our analysis expanded over 10-fold the number of ubiquitinated proteins identified under arsenite stress (35, 44, 45) (**Fig. 2C**), highlighting the power of our DIA and subcellular fractionation method to providing a deep resource of localized ubiquitin signals under stress. These enriched peptides significantly correspond to proteins involved in various immune signaling pathways related to bacterial and viral infections, as well as translation regulation through ribosome biogenesis and tRNA-aminoacyl biosynthesis (**Fig. 2D**). Consistent with other studies (35, 46), most proteins had three or fewer stress-enriched ubiquitin sites (**Fig. 2E**), suggesting ubiquitination during stress is distributed across many targets, rather than high levels of ubiquitin on relatively few targets. Among targeted proteins, GCN1 and ribosomal proteins had high numbers of unique non-cytosolic ubiquitin peptides enriched under stress (**Fig. 2E, supplemental Table S2**), suggesting ubiquitination might be involved in localized translation control (47). Supporting our previous results that subcellular K63ub accumulation occurs locally (**Fig. 1H**), we also found that increases in ubiquitinated peptides are not determined by localized increases in protein abundance (**Fig. 2F**). These proteomic analyses not only mapped the subcellular landscape of ubiquitin signals but also highlighted how prominent localized ubiquitin dynamics are under arsenite stress. Our findings suggest a massive rewiring of a broad range of signaling pathways involved in stress response, cellular physiology, and quality control mechanisms.

**Figure 2.**
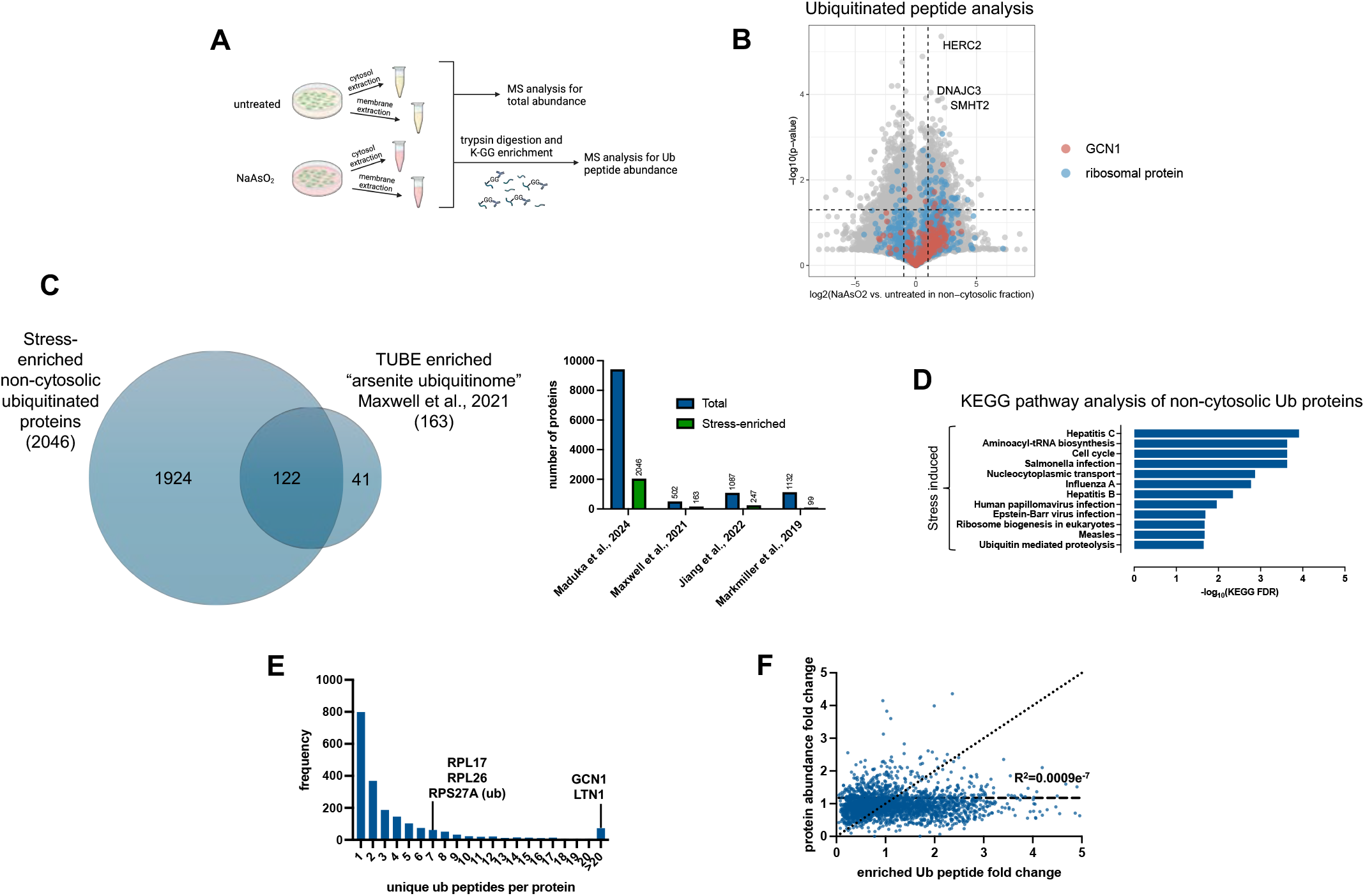
Ubiquitin targets non-cytosolic stress response factors in response to arsenite. 2A – Schematic of sample preparation for ubiquitin peptide proteomic analysis during arsenite stress. 2B – Volcano plots showing enrichments under arsenite stress of ubiquitin peptides in the non-cytosolic fractions. Peptides for GCN1 and various ribosomal proteins are highlighted, as well as top ubiquitinated peptides. 2C – Venn diagram and coverage of total proteins and those enriched under arsenite stress from ubiquitin proteomics dataset compared to Maxwell et al. 2021, Jiang et al. 2023, and Markmillan et al. 2019. 2D – KEGG pathway enrichment of proteins from ubiquitin peptides enriched in the non-cytosolic fraction under arsenite stress (fold change > 3 and q-value < 0.05). 2E – Frequency of proteins by the number of unique di-glycine peptides detected in the non-cytosolic fraction under stress. 2F – Correlation plot of fold changes of ubiquitin peptides with the fold change of the abundance of their corresponding proteins.

Because K63ub has non-proteolytic functions (37), we next sought to identify factors that respond to accumulated ubiquitin for downstream processing during arsenite stress. To do this, we performed a new set of proteomic analyses in cytosolic and non-cytosolic compartments of cells treated with or without NaAsO_2_ after enrichment of the compartments where K63ub chains accumulate (**Fig. 3A**). To better understand the precision of our fractionation method, we first quantitatively accounted for the presence of organellar proteins in the non-cytosolic fraction. Indeed, we found many membrane-annotated proteins, such as those localized to the ER, mitochondria, and endosomes, enriched in non-cytosolic purified compartments (**Fig. 3B**), with the most abundant of proteins in these compartments being ribosomal proteins likely associated with the ER (**supplemental Fig. S5A**). Amongst all fractions and treatments, we identified 5,175 unique proteins associated to these purified non-cytosolic compartments and determined that 1,434 proteins were significantly enriched in comparison to the cytosol (**supplemental Table S3**). Supporting previous data (**supplemental Fig. S3A-S3C**), UBE2A and UBE2N, known to be involved in K63ub signaling, were not enriched in the non-cytosolic fraction under stress (**supplemental Fig. S3D**). As our dataset characterized the landscape of proteins that are significantly enriched in non-cytosolic compartments, further analyses could reveal proteins that associate with K63ub chains, providing insights in the downstream pathways mediated by these signals during stress. Through gene ontology analysis, we found that 64 proteins in our whole dataset had ubiquitin binding functions, and 31 of these were found to be significantly enriched (over 1.5-fold) during arsenite stress in the non-cytosolic compartment (**Fig. 3C, supplemental Table S3**). These enriched proteins with UBDs are involved in processes such as NF_*k*_B signaling and defense response to bacterium (**Fig. 3E**), which is consistent both with the ubiquitin peptide enrichment (**Fig. 2D**) and with functions of K63ub signaling described in the literature (48). Interestingly, among the enriched proteins with UBDs were many adaptors of the ubiquitin processing protein VCP (**Fig. 3C-D**). VCP is a AAA+ ATPase known to associate and process various ubiquitinated substrates involved in pathways such as proteasomal degradation, autophagy, ribosome quality control, and stress granule disassembly (49–51). To this function, VCP is assisted by a variety of adaptors that bind and recognize different polyubiquitin chains in different cellular compartments (52). Over 60% of these VCP adaptors were uniquely enriched in the non-cytosolic compartment, suggesting they may be involved in local ubiquitin signaling. Furthermore, FAF2, UBXN1, and NPLOC4, which are among the significantly enriched proteins, have been described in various stress responses within the ER and mitochondria (27, 53, 54). These data indicate that the VCP system may participate in localized ubiquitin signaling following arsenite stress.

**Figure 3.**
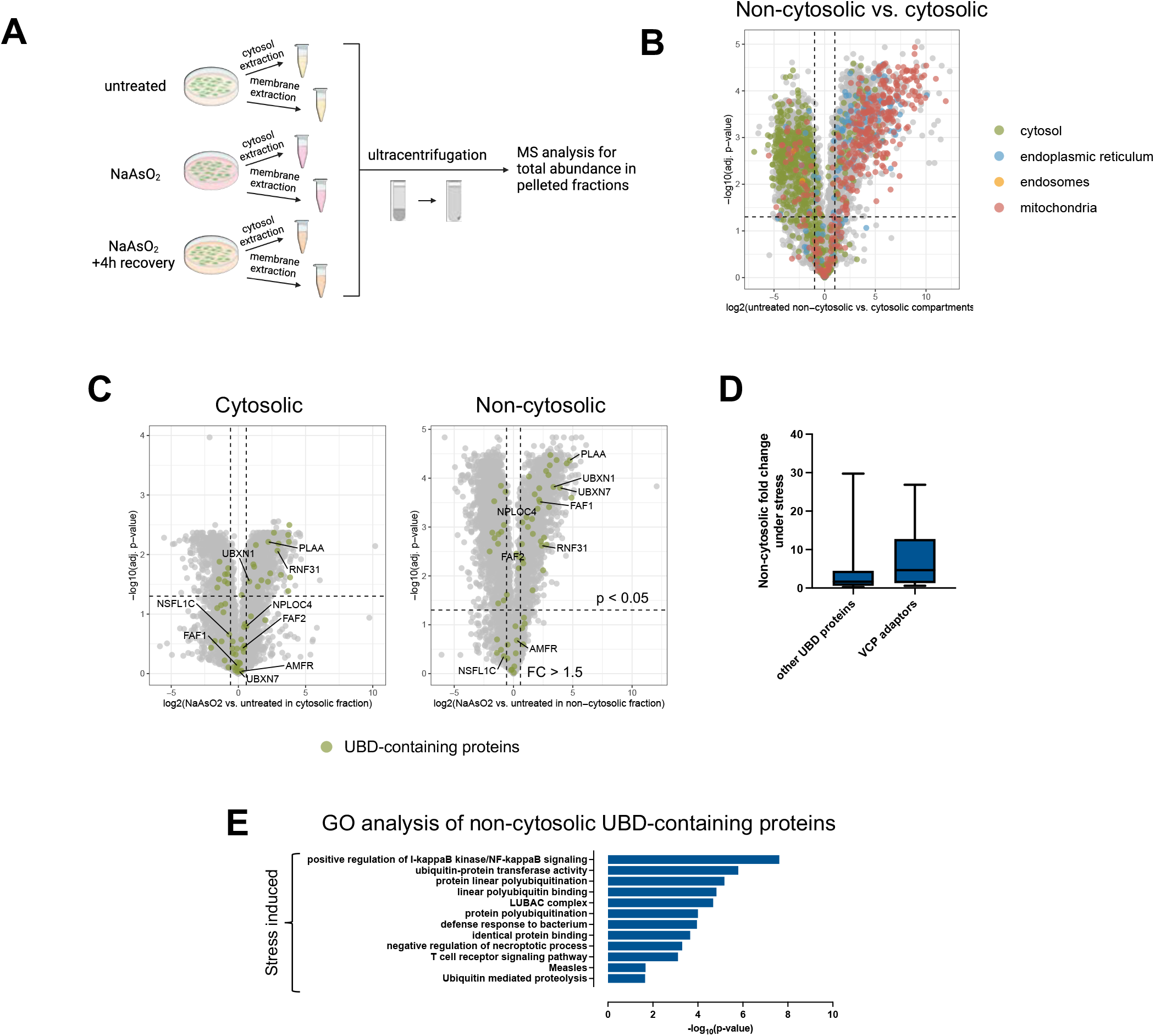
Various ubiquitin processors are enriched in purified non-cytosolic fractions in response to arsenite. 3A – Schematic of sample preparation for proteomics analysis of the cytosolic and non-cytosolic fractions during arsenite stress. 3B – Volcano plot of log_2_(fold enrichment) of various proteins in the non-cytosolic versus cytosol fractions. Highlighted proteins are localized to the cytosol, endoplasmic reticulum, mitochondria, and endosomes. Lists were generated from The Human Protein Atlas. 3C – Volcano plots showing enrichments under arsenite stress of proteins with ubiquitin binding domains in the non-cytosolic and cytosolic fractions. Known VCP adaptors were highlighted among the proteins with UBDs in both plots. 3D – Fold change under arsenite stress of VCP adaptors and other identified UBD-containing proteins within the non-cytosolic fraction. 3E – Gene ontology enrichment of proteins with ubiquitin binding domains enriched in the purified non-cytosolic fraction under arsenite stress (fold change > 1.5 and p-value < 0.05).

### Arsenite stress induces ubiquitin-mediated subcellular partitioning of VCP

As many of these adaptors of VCP were enriched in the non-cytosolic compartments under stress in our mass spectrometry dataset (**Fig. 3C**), we hypothesized that VCP is also partitioned to this compartment during oxidative stress. With an antibody against endogenous VCP, we first found that VCP accumulates more in the non-cytosolic fraction during arsenite stress compared to the cytosolic fraction (**Fig. 4A**). We confirmed this finding by showing that exogenously expressed FLAG-VCP also accumulates in non-cytosolic compartments under arsenite stress (**supplemental Fig. S6A**). To test whether the recruitment of VCP is mediated by ubiquitin, we prevented ubiquitination by preincubating cells with the E1 inhibitor TAK-243 (**Fig. 4B**). Inhibition of ubiquitination prevented VCP accumulation in the non-cytosolic fraction under arsenite stress (**Fig. 4C**), suggesting that VCP recruitment is not a product of stress itself but relies on the increase of ubiquitin chains. As we previously showed that the levels of K63ub chains are reversible (**Fig. 1F**), we tested whether VCP accumulation also reverses in stress recovery. We found that VCP accumulation is reversible and correlates with the levels of ROS and with the reversal of K63ub signals (**Fig. 4D-E**). As VCP recruitment is dependent on ubiquitination and is reversed during arsenite recovery, we asked whether VCP associates with non-cytosolic K63ub. Using confocal microscopy, we found that K63ub puncta colocalizes with VCP with increasing overlap during stress (**Fig. 4F**), suggesting that K63ub signals might be processed by VCP. Lastly, VCP accumulation in non-cytosolic compartments appears to be stress-specific, and not be solely mediated by other stress signals upstream of ubiquitination events such as ER stress and stress granule formation (**supplemental Fig. S6B-S6C**). Together, our results suggest that VCP partitions to non-cytosolic compartments in a process mediated by K63ub chains that accumulate under arsenite stress.

**Figure 4.**
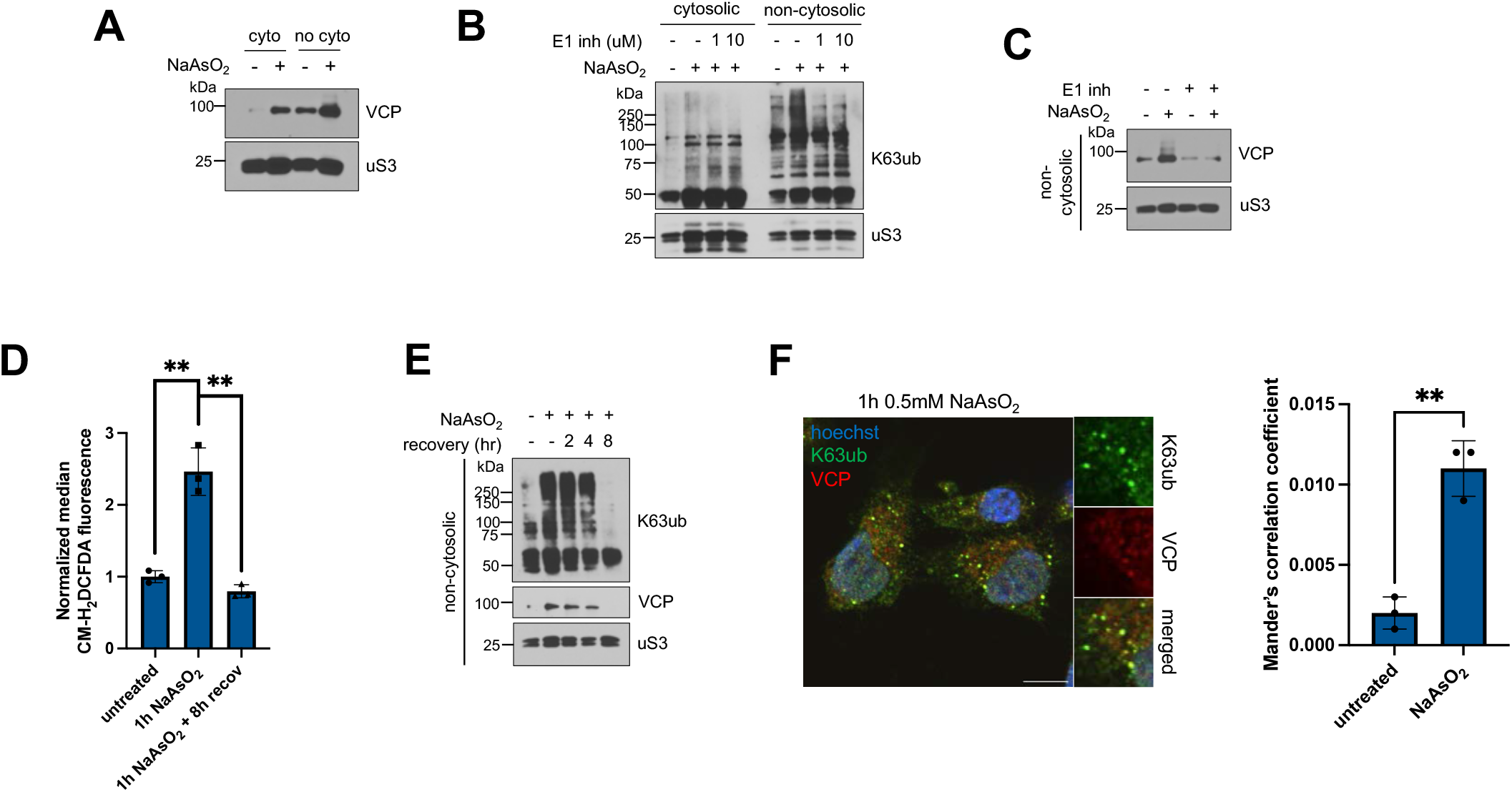
Arsenite stress promotes ubiquitin-dependent non-cytosolic VCP accumulation. 4A – Arsenite stress induces VCP accumulation in purified non-cytosolic compartments. Immunoblots of anti-VCP from fractions of cells treated with 1 hour of 0.5 mM NaAsO_2_. Anti-uS3 was used as a loading control for each purified fraction. 4B – Immunoblots of anti-ubiquitin from fractions of cells, and anti-K63ub from fractions treated with 1 hour of 0.5 mM NaAsO_2_ without or with increasing concentrations of the E1 inhibitor TAK-243. Anti-uS3 was used as a loading control for each purified fraction. 4C – E1 inhibition prevents arsenite-induced VCP accumulation. Immunoblot of anti-VCP from non-cytosolic fractions of cells preincubated without or with 1 μM of the E1 inhibitor TAK-243 for 3 hours, then treated with 1 hour of 0.5 mM NaAsO_2_. Anti-uS3 was used as a loading control. 4D – Arsenite stress induces ROS accumulation. Cells were either untreated, treated with 1 hour of 0.5mM NaAsO_2_, or allowed to recover, then stained with the ROS agent CM-H2DCFDA. Levels of ROS per cell were processed using flow cytometry. 4E – VCP accumulation is reversible after arsenite washout and follows dynamics of K63ub reversal. Immunoblots of anti-K63ub and anti-VCP from non-cytosolic fractions of cells along a course of treatment with 1 hour of 0.5 mM NaAsO_2_ with a washout and recovery of either 2, 4 or 8 hours. Anti-uS3 was used as a loading control. 4F – K63ub puncta present during arsenite stress colocalizes with VCP. Immunofluorescence microscopy of anti-K63ub and anti-VCP from cells treated with 1 hour of 0.5 mM NaAsO_2_. Cells were prepared with a digitonin wash to release cytosolic components before fixation. Hoechst was used to identify cells. Scale bar = 10 *μ*m. Manders correlation coefficient was calculated to determine percent occupancy of K63ub puncta in VCP signal.

It is known that VCP recruitment to ubiquitinated substrates is mediated by a family of ubiquitin binding adaptors (49). As many VCP adaptors were enriched in the non-cytosolic compartment under stress (**Fig. 3C-D**), we sought to identify the adaptors involved in VCP subcellular partitioning. We first tested whether FAF2, a known ER-localized VCP adaptor demonstrated to associate with K63ub linkages (27), aids in local VCP accumulation during arsenite stress. However, depletion of FAF2 levels by siRNA depletion did not impair VCP accumulation (**supplemental Fig. S6D**). We then returned to our proteomics dataset and found that the VCP adaptor NPLOC4 was enriched in the non-cytosolic fraction under stress compared to steady-state conditions (**Fig. 3C**). Importantly, NPLOC4 was also only enriched in non-cytosolic compartments under stress with a 2.1-fold change (**Fig. 3C, supplemental Table S3**), implying a local response and highlighting it as a strong candidate for further analysis. NPLOC4 is an ER-localized adaptor facilitating VCP’s involvement in ERAD, RQC, and other pathways through binding K63ub and K48ub chains (55). We observed that NPLOC4 localizes to non-cytosolic compartments by western blot as well (**Fig. 5A**). To explore the role of NPLOC4 in VCP partitioning to the non-cytosolic compartment during stress, we pre-incubated cells with the NPLOC4 inhibitor disulfiram (56, 57), and found VCP accumulation was decreased (**Fig. 5B**). We then depleted NPLOC4 with siRNA and found that VCP accumulation to the non-cytosolic fraction during arsenite stress was also decreased (**Fig. 5C**), both supporting NPLOC4 involvement in VCP recruitment to non-cytosolic compartments. The data supports that NPLOC4 is partially supporting a ubiquitin-mediated partitioning of VCP to non-cytosolic compartments under arsenite stress.

**Figure 5.**
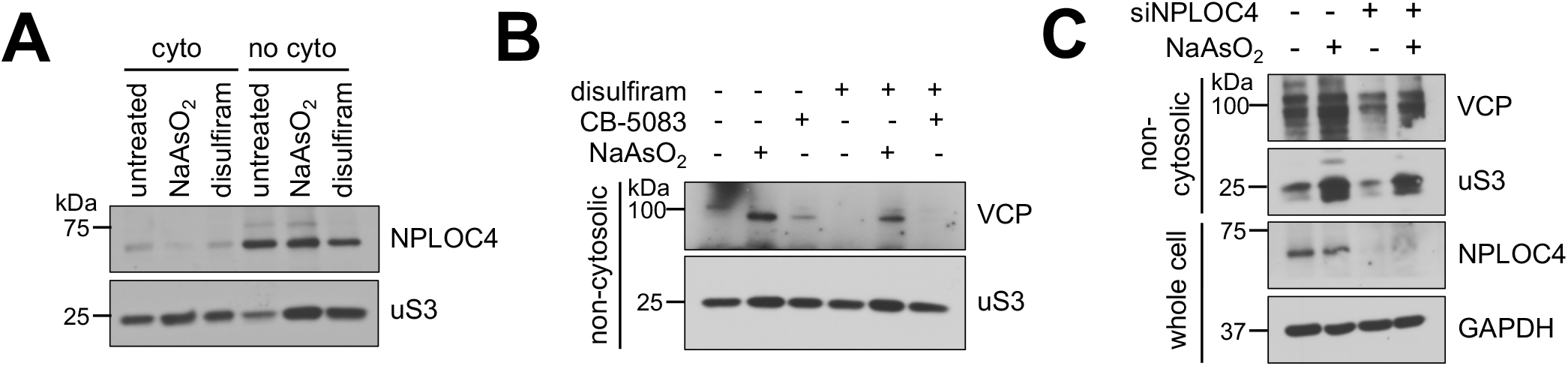
VCP adaptor NPLOC4 assists in arsenite-induced non-cytosolic VCP accumulation. 5A – NPLOC4 localizes to the non-cytosolic compartments. Immunoblots of anti-NPLOC4 from fractions of cells treated with 1 hour of 0.5 mM NaAsO_2_ or 5 hours of 125 μM disulfiram. Anti-uS3 was used as a loading control for each purified fraction. 5B – NPLOC4 inhibition with disulfiram reduces arsenite-induced VCP accumulation in purified non-cytosolic fractions. Immunoblots of anti-VCP from non-cytosolic fractions of cells treated with DMSO or 125 μM of the NPLOC4 inhibitor disulfiram for 4 hours, then without or with 1 hour of 0.5 mM NaAsO_2_. Anti-uS3 was used as a loading control for each purified fraction. 5C – NPLOC4 knockdown reduces arsenite-induced VCP accumulation. Immunoblots of anti-VCP from non-cytosolic fractions of cells transfected with siVCP or siControl, then treated without or with 1 hour of 0.5 mM NaAsO_2_. Anti-uS3 was used as a loading control. Successful knockdown of NPLOC4 protein levels were assessed from total cell lysates with anti-NPLOC4, using anti-GAPDH as a loading control.

### VCP participates in recycling of non-cytosolic K63 ubiquitin signals

Our findings show that VCP is recruited to the non-cytosolic compartments through its association with K63ub linkages. Thus, this model implies that ubiquitination happens upstream of VCP recruitment, and the total K63ub accumulated is defined by a balance of ubiquitin conjugation and VCP processing. Disrupting the VCP system is known to lead to accumulation of ubiquitin conjugates involved in both proteasome-dependent and independent signaling (19, 58). Therefore, we tested whether inhibition of VCP would lead to accumulation of K63ub even in the absence of stress. We found that CB-5083, an inhibitor of VCP’s D2 ATPase domain (59, 60), also causes accumulation of K63ub specifically in non-cytosolic compartments (**Fig. 6A**). We corroborated this finding by depleting VCP with siRNA, finding that K63ub also accumulates in the non-cytosolic fraction (**Fig. 6B**). Correlating with these results, CB-5083 increased non-cytosolic K63ub puncta (**Fig. 6C**), corroborating that VCP is indeed involved in processing K63ub chains that accumulate under arsenite stress. Interestingly, we observed that VCP itself under CB-5083 treatment accumulated in the non-cytosolic fraction (**Fig. 6A**). This implies that impairing VCP processing and not recruitment independently leads to VCP buildup in a feedforward regulatory mechanism, recruiting more VCP needed for ubiquitin chain processing. Since NPLOC4 seems to be involved in VCP recruitment during arsenite stress (**Fig. 5**), we tested whether NPLOC4 inhibition would also lead to K63ub accumulation. Inhibition of NPLOC4 with disulfiram similarly led to localized K63ub accumulation (**Fig. 6D**) as well as K63ub puncta formation (**Fig. 6E**), both suggesting the recruitment of VCP mediated by NPLOC4 is important for recycling K63ub. Furthermore, we found that non-cytosolic K63ub puncta do not respond with an additive effect when combining CB-5083 and arsenite stress (**Fig. 6F-H**), reinforcing that VCP is involved in K63ub accumulation.

**Figure 6.**
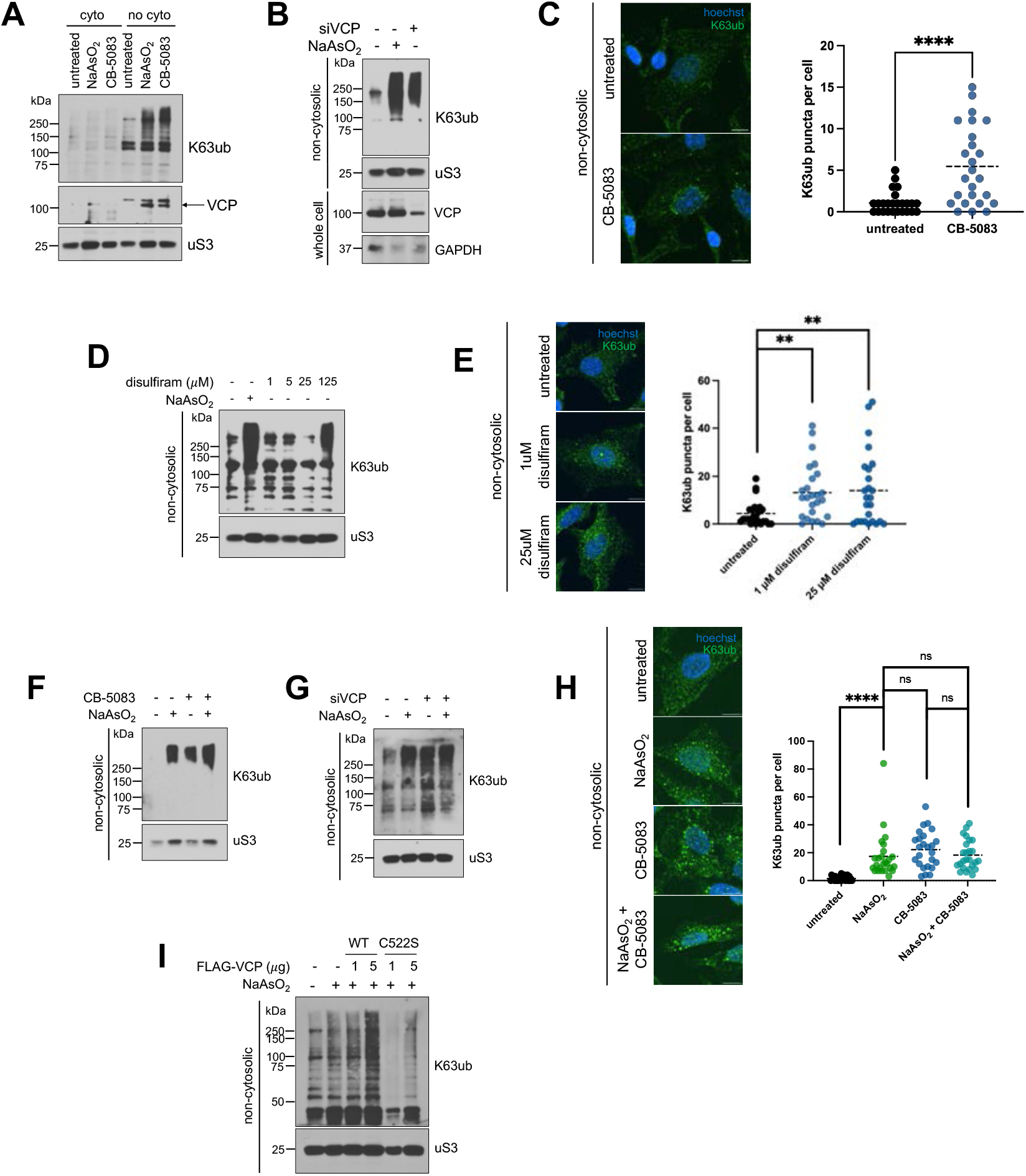
Disruption of VCP system function leads to non-cytosolic K63ub accumulation. 6A – VCP inhibition with CB-5083 induces local accumulation of K63ub and VCP in purified non-cytosolic fractions. Immunoblots of anti-K63ub, anti-K48ub, and anti-VCP from fractions of cells treated with either 1 hour of 0.5 mM NaAsO_2_ or 4 hours of 5 μM of the VCP inhibitor CB-5083. Anti-uS3 was used as a loading control for each purified fraction. 6B – VCP knockdown promotes K63ub accumulation. Immunoblots of anti-K63ub from non-cytosolic fractions of cells treated with either 1 hour of 0.5 mM NaAsO_2_ or transfected with siVCP versus siControl. Anti-uS3 was used as a loading control. Successful knockdown of VCP protein levels were assessed from total cell lysates with anti-VCP, using anti-GAPDH as a loading control. 6C – Non-cytosolic K63ub chains form puncta upon VCP inhibition. Immunofluorescence microscopy of anti-K63ub from cells treated with either 1 hour of 0.5mM NaAsO_2_, 4 hours of 5 μM CB-5083, or both. Cells were prepared with a digitonin wash to release cytosolic components before fixation. Hoechst was used to identify and count cells. Analysis included 25 cells per group. Scale bar = 10 *μ*m. 6D – Inhibition of NPLOC4 with disulfiram induces K63ub accumulation in purified non-cytosolic fractions. Immunoblots of anti-K63ub from non-cytosolic fractions of cells treated with either 1 hour of 0.5mM NaAsO_2_ or increasing concentrations of disulfiram. Anti-uS3 was used as a loading control. 6E – Non-cytosolic K63ub linkages form puncta upon NPLOC4 inhibition. Immunofluorescence microscopy of anti-K63ub from cells treated with either 1 μM or 25 μM disulfiram. Cells were prepared with a digitonin wash to release cytosolic components before fixation. Hoechst was used to identify and count cells. Analysis included 25 cells per group. Scale bar = 10 *μ*m. 6F – Combining arsenite stress and VCP inhibition does not induce additive K63ub accumulation. Immunoblots of anti-K63ub, anti-K48ub, and anti-VCP from non-cytosolic fractions of cells treated with either 1 hour of 0.5 mM NaAsO_2_, 4 hours of 5 μM CB-5083, or both. Anti-uS3 was used as a loading control. 6G – Combining arsenite stress and VCP knockdown does not induce additive K63ub accumulation. Immunoblots of anti-K63ub from non-cytosolic fractions of cells treated with either 1 hour of 0.5 mM NaAsO_2_, transfected with siVCP versus siControl, or both. Anti-uS3 was used as a loading control. 6H – Non-cytosolic K63ub signal induction is not additive when combining CB-5083 and arsenite stress. Immunofluorescence microscopy of anti-K63ub from cells treated with either 1 hour of 0.5mM NaAsO_2_, 4 hours of 5 μM CB-5083, or both. Cells were prepared with a digitonin wash to release cytosolic components before fixation. Hoechst was used to identify and count cells. Analysis included 25 cells per group. Scale bar = 10 *μ*m. 6I – Overexpression of VCP decreases levels of accumulated K63ub during arsenite stress. Immunoblots of anti-K63ub from non-cytosolic fractions of cells transfected with increasing amounts of FLAG-VCP WT or FLAG-VCP C522S plasmids for 48 hours, then treated the following day with 1 hour of 0.5 mM NaAsO_2_. Anti-uS3 was used as a loading control for each purified fraction. Successful transfections of FLAG-VCP were assessed under total cell lysate with anti-FLAG, using anti-GAPDH as a loading control.

**Figure 7.**
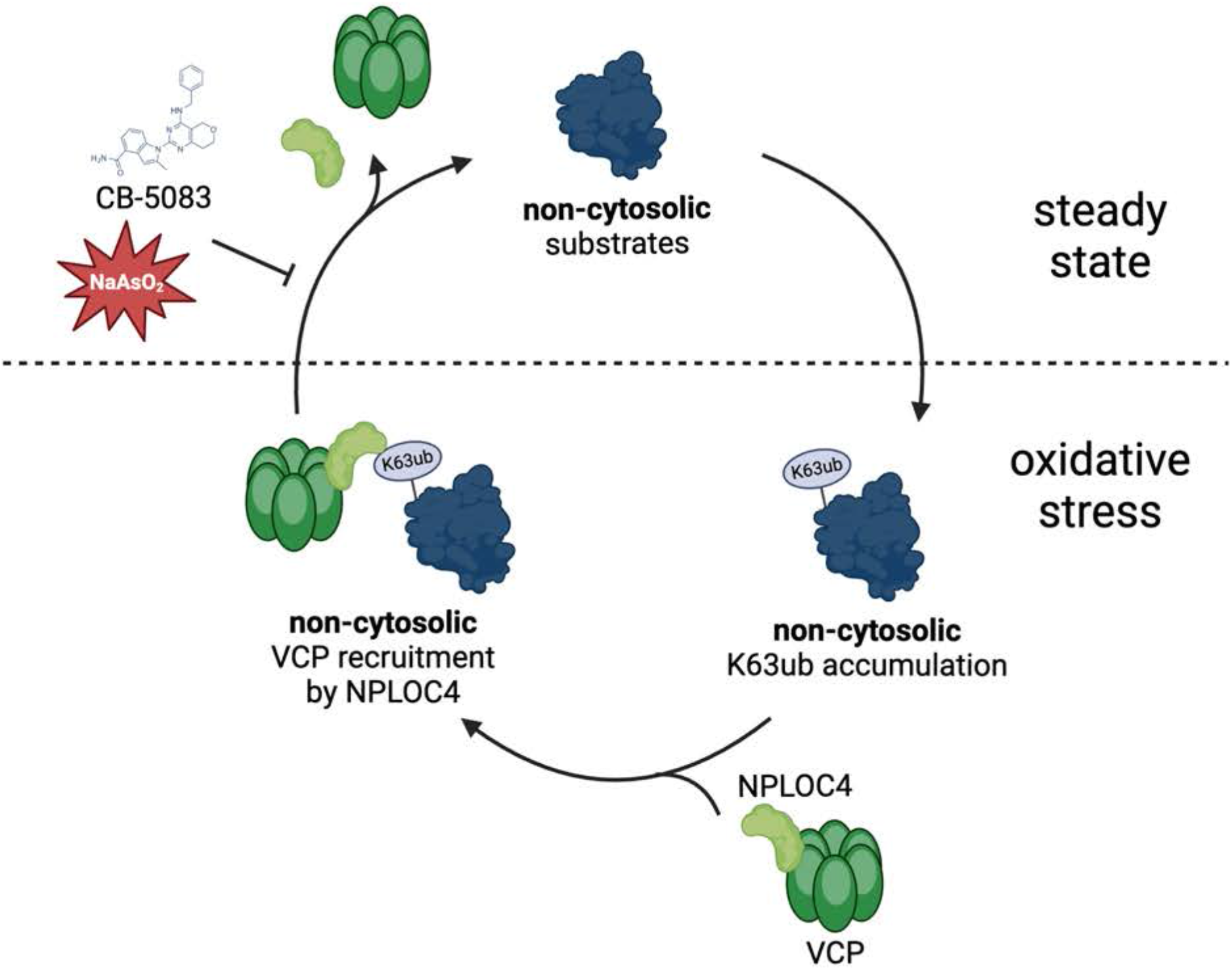
Model depicting relationship between non-cytosolic K63 ubiquitin accumulation and VCP regulation under stress.

Our results suggest a model in which a cyclical accumulation and reversal of K63ub is dependent on VCP activity (**Fig. 6A**). The accumulation of K63ub chains upon arsenite exposure led us to hypothesize that ROS generated by arsenite could be directly inhibiting VCP activity, which would cause the buildup of K63ub chains. VCP processing activity has been shown to be sensitive to oxidative stress by diamide, so we next asked whether arsenite stress is directly affecting VCP function. It was previously shown that VCP cysteine 522 (C522) residue can be oxidized under stress and that mutant VCP (C522S) is resistant to redox inhibition (61). In this context, expression of the stress-resistant mutant VCP^C522S^ would lead to continuous processing and a reduction in the levels of K63ub chains that accumulate under stress. Therefore, we tested whether VCP’s redox regulation via C522 could be underlying the accumulation of K63ub chains during arsenite stress. Indeed, we found that expression of VCP^C522S^ but not VCP^WT^ led to low levels of K63ub chains during arsenite stress (**Fig. 6I**), suggesting oxidation from arsenite may regulate VCP’s ability to process K63ub. Altogether, our data indicate that redox regulation of VCP activity mediates the recycling of non-cytosolic K63ub signals during oxidative stress.

## DISCUSSION

Ubiquitination is a critical signaling mechanism for protection under environmental stress, and cells have adapted intricate responses utilizing ubiquitin signals that depends on the context of the stressor. We have shown that upon oxidative stress, K63ub signals accumulate in non-cytosolic compartments (**Fig. 1**) and utilized K-GG remnant ubiquitin proteomics to dissect the various pathways involving non-cytosolic ubiquitination events during oxidative stress (**Fig. 2**). We show many of these localized ubiquitination events are connected to defense during various viral and bacterial infections, as well as the control of protein synthesis, which are known to be highly dynamic and regulated under stress conditions (9, 62). This work revealed that unique combinations of biochemical, cellular, and proteomics approaches has the power to uncover underlying regulatory signals that help us dissect the complexity and dynamics of PTMs during the stress response. As we continue to elucidate the rules of subcellular ubiquitin signaling under stress, chain specific isolation approaches such as TUBE enrichment (63) can be combined with mass spectrometry analysis to fully understand participation of additional ubiquitin linkages beyond K63ub in the stress response. Additionally, further subcellular fractionation will increase resolution and provide further details on new stress response mechanisms mediated by K63ub signals under stress.

Many groups have recently uncovered mechanisms involving non-canonical ubiquitin signaling within or associated to various organelles like the mitochondria, ER, and endosomes (24, 25, 64). Our work highlighted that K63ub signals partition to non-cytosolic compartments potentially modifying a large fraction of proteins from different organelles and membrane components (**supplemental Fig. S2-S3, Fig. 2-3**). Our work generated a high-resolution proteomics resource adding a series of new elements to the pool of ubiquitin substrates regulated in a subcellular fashion under conditions of stress (**supplemental Table S2, Fig. 2**). Based on the high number of proteins that we identified to be differentially ubiquitinated under stress (**Fig. 2E-F**), it would be surprising that only a few selected E2 ubiquitin conjugases and E3 ligases would account for this abundant and widespread modification. The human genome encodes about 40 E2s and over 600 E3s distributed across all cellular compartments (20), which offer diversity of roles and define substrate and linkage specificity (18). However, we established that regulation of K63ub processing particularly by the VCP system is a main driver of this subcellular ubiquitin accumulation (**Fig. 6**). These findings suggest a model in which several ubiquitin enzymes are important for its conjugation, which are sorted by VCP in partnership with its many adaptors including NPLOC4 (**Fig. 5**). These events could be contributing to a rewiring of cellular machinery towards defense against oxidative stress, and elucidation of the downstream pathways triggered by VCP will deepen our understanding of localized and global stress responses.

VCP regulation is evident in multiple contexts during steady-state and during responses to environmental stimuli. For example, VCP can be regulated through oxidation that impairs its activity and its association to its adaptors (61). Other non-ubiquitin binding VCP adaptors like VCF1 have been shown to promote VCP/NPLOC4/UFD1 association to its ubiquitinated substrates (65). We have shown that upon oxidative stimuli, VCP is regulated by subcellular localization and the majority is partitioned in non-cytosolic compartments (**Fig. 4**). Our data further suggests ROS produced by sodium arsenite directly leads to impaired VCP activity, as expression of oxidation-resistant mutant VCP^C522S^ prevents K63ub accumulation under stress (**Fig. 6I**). Functionally, VCP is involved in many ubiquitin-dependent stress mechanisms such as ribosome quality control and mitophagy, in which VCP degrades ubiquitinated nascent peptides upon aberrant translation or ubiquitinated proteins in the outer mitochondrial membrane, respectively (66, 67). Our proteomic analysis showed several ribosomal and mitochondrial proteins to be ubiquitinated under stress, which corroborates that supporting organellar quality and function is a key role of the VCP system under steady state conditions. VCP redox inhibition might support a rapid regulatory mechanism by which cells rewire their energetic use (49) to favor pressing stress-related pathways. After stress cessation, reversal of these ubiquitin signals occurs with participation of VCP reactivation (**Fig. 4E**), which is critical to restore cellular proteostasis, quality control pathways, and growth.

VCP processes ubiquitinated signals for both degradation and non-degradative mechanisms (68). Disruptions of these functions of VCP are linked to diseases, such as inclusion body myopathy, Paget disease of bone and frontotemporal dementia (69), emphasizing its importance in cellular physiology. Additionally, inhibitors of VCP such as CB-5083 and other derivatives have reached clinical trials for its anti-tumor activity (70). Our data shows that VCP follows a subcellular pattern under oxidative stress to that of proteasome-independent K63ub signals (**Fig. 4-6**). Our findings indicate localized increases in VCP-K63ub signaling may serve as a response to oxidative stress in many disease contexts that can be therapeutically targeted to increase cellular sensitivity in diseases like cancer and infection (62, 71). Overlapping connections of VCP and K63ub signaling to diseases highlight importance of our described localized stress signaling mechanism. As our work broadens the potential roles of VCP, co-targeting immune signaling or translation control pathways in the clinic could potentialize anti-tumor therapies and provide synergistic therapeutic effects to fight malignancy. This work has potential to highlight avenues for intervention of VCP and noncanonical ubiquitin signaling in diseases involving oxidative stress.

## MATERIALS AND METHODS

### Mammalian cell culture and chemical treatments

HeLa, U2OS, and HEK293T cell lines were all obtained from ATCC. HeLa cells were grown in EMEM (ATCC, 30-2003), U2OS cells grown in McCoy’s 5A (ATCC, 30-2007), and HEK293T cells grown in DMEM (ATCC, 30-2003). All cell lines were grown in media supplemented with 10% fetal bovine serum and 1% penicillin/streptomycin, and incubated at 37C and 5% CO_2_. Cells were confirmed free of mycoplasma infection quarterly. For oxidative stress induction, cells were treated with 0.5 mM sodium arsenite (Sigma, S7400) diluted in PBS, for one hour before harvesting. For VCP inhibition, cells were treated with 5 μM CB-5083 (Selleck, S8101) for 4 hours before harvesting. For NPLOC4 inhibition, cells were treated with disulfiram (Tocris, 3807) unless otherwise stated.

### Sequential detergent fractionation

Non-cytosolic and cytosolic fractionations were performed typically on ice with cells at 80% confluence. First, microtubules were depolymerized by incubating cells on ice with ice cold PBS for 10 minutes. Then cytosolic extraction buffer (110 mM KCl, 15 mM MgCl_2_, 4 mM CaCl_2_, 25 mM HEPES, 0.03% digitonin, EDTA-free protease inhibitor tablet (Roche, 04693159001), 20 mM iodoacetamide) was added to cells to cover the plate for 15 minutes. Then cytosolic wash buffer (110 mM KCl, 15 mM MgCl_2_, 4 mM CaCl_2_, 25 mM HEPES, 0.006% digitonin) was added for 5 minutes to remove residual cytosolic components. Then DDM lysis buffer (200 mM KCl, 15 mM MgCl_2_, 4 mM CaCl_2_, 25 mM HEPES, 2% DDM, protease inhibitor tablet, 20 mM iodoacetamide) was added to cells for 15 minutes. Both components were then centrifuged at 12,700 rpm for 15 minutes to remove cellular debris.

### Whole cell lysis

After indicated treatments, cells were washed and harvested by trypsinization. Then, cells were resuspended on ice with RIPA lysis buffer (150 mM Tris-HCl pH 7.4, 150 mM NaCl, 1 mM EDTA, 1% IGEPAL, 0.1% SDS, protease inhibitor cocktail), and rotated at 4C for 30 minutes. Samples were then cleared by centrifugation at 12,700 rpm for 15 minutes, and protein concentrations were determined by Bradford assay (Bio-Rad, 500-0205) and normalized prior to western blotting.

### siRNA and plasmid transfections

For RNAi knockdowns, cells were transfected at about 70% confluency with either siGENOME non-targeting control siRNA (Dharmacon, D-001206-13-05) or indicated siGENOME SMARTpool siRNA (Dharmacon) using Lipofectamine RNAiMAX (ThermoFisher, 13778100) according to the manufacturer’s instructions. The following siRNAs were used: siRNA control pool (D-001206-13), siNPLOC4 (M-020796-01), siVCP (M-008727-01), siG3BP1 (M-012099-02), siUBE2A (M-009424-00), siUBE2B (M-009930-00), siUBE2N (M-003920-01), siFAF2 (M-010649-01). For plasmids, cells were transfected using Lipofectamine 3000 (ThermoFisher, L3000001) according to the manufacturer’s instructions. After 24 hours, transfection mix was removed and cells were resuspended then seeded equally between planned treatment conditions in both 10-cm and 6-well plates. All cells were then grown for 24-48 hours, and cells in 6-well plates were underwent total cell lysis to assess knockdown efficiency or protein expression by western blot, while cells in 10-cm plates underwent sequential detergent fractionation or other intended assay.

### Immunoblotting

Whole lysate, fractions, or purified fractions were boiled in Laemmli buffer were loaded onto a 10-15% SDS-PAGE gel and resolved, transferred onto a PVDF membrane, and blocked with 5% BSA in TBS-T. Membranes were incubated with primary antibody for one hour and secondary antibodies, both in 5% BSA in TBST. The following primary antibodies and concentrations were used: anti-K63 ubiquitin (1:5,000, EMD Millipore, 051308), anti-K48 ubiquitin (1:5,000, Cell Signaling Technology, 8081S), anti-RPS3/uS3 (1:10,000, Cell Signaling, #9538S), anti-tubulin (1:7,500, Novus Biologicals, MAB93441), anti-TRAPa (1:10,000, Abcam, ab240562), anti-puromycin (1:7,500, MilliporeSigma, MABE343), anti-VCP (1:2,500, Santa Cruz Biotechnology, #sc-57492), anti-NPLOC4 (Proteintech, 11638-1-AP), anti-GAPDH (1:10,000, Abcam, ab9485), anti-FLAG (1:2,500, MilliporeSigma, C986X12), anti-ubiquitin/P4D1 (1:5,000, Cell Signaling, 3936S), anti-BiP/GRP78 (1:2,500, Invitrogen, MA5-27686), anti-G3BP (1:2,500, BD Biosciences, 611126), anti-Rad6/UBE2A (1:2,500, Abcam, ab31917), anti-UBE2N (1:2,500, Cell Signaling, 6999S), anti-FAF2 (1:2,500, Proteintech, 16251-1-AP). Both anti-Rabbit IgG HRP (Cytiva, NA934), and anti-Mouse IgG HRP (Cytiva, NA931) were used as secondary antibodies. Densitometry for western blot quantification was performed using FIJI software.

### Sucrose sedimentations

Total cell lysates, cytosolic fractions, or non-cytosolic fractions were loaded over a 50% sucrose solution (in 50 mM Tris-acetate pH 7, 150 mM NaCl and 15 mM MgCl_2_) and ultracentrifuged for 2 hours at 70,000 rpm in a Beckman Optima Max-TL with a TLA-110 rotor. Pellets were resuspended in 50 mM Tris-acetate, pH 7, 150 mM NaCl. Concentrations were measured by NanoDrop for rRNA as an approximation for ribosome content, normalized, then samples were used for intended assays

### Immunofluorescence microscopy

HeLa cells were plated on coverslips in 6-well plates at 250,000 cells per well and allowed to grow one day to about 80% confluency. After indicated treatments, cells were washed with ice-cold PBS for 10 minutes. Then cytosolic extraction buffer (110 mM KCl, 15 mM MgCl_2_, 4 mM CaCl_2_, 25 mM HEPES, 0.03% digitonin) was added to the coverslips in the wells for 15 minutes. Then cells were permeabilized and fixed with ice-cold 100% methanol. Cells were then stained with anti-K63 ubiquitin (1:1,000, EMD Millipore, 051308), and occasionally anti-VCP (1:500, BD Biosciences, 612182) or anti-G3BP (1:2,000, BD Biosciences, 611126), all diluted in 5% BSA in TBS-T. Afterwards, cells were stained with anti-rabbit Alexa Fluor 488 (1:1,000, Abcam, ab150077), and occasionally anti-mouse Alexa Fluor 647 (1:1,000, Abcam, ab150115) diluted in 5% BSA in TBS-T. Nuclei were then stained with Hoechst dye (1:1,000, Thermo, 62249). Coverslips were mounted using Anti-Fade Fluorescence mounting medium (Abcam, ab104135), sealed on slides and imaged either on an Andor Dragonfly Spinning Disk confocal microscope (with iXon EMCCD camera and controlled by FUSION software) at 100x/1.40-0.70, or a Leica Thunder Imager (with sCMOS camera and controlled with LAS X software) at 63x magnification. The following excitation and emission parameters were used: 488/519 for Alexa Fluor 488, 637/668 for Alexa Fluor 647, 405/ for Hoechst. All images were analyzed using FIJI software. K63ub channels were processed using the gaussian blur filter. Then, thresholding was used on the K63ub channel to indicate K63ub puncta, starting with NaAsO_2_-treated samples. Regions of interest were drawn around 25 cells per image, and number of puncta were determined by counting number of K63ub particles above the threshold. Colocalization was determined by generating Mander’s correlation coefficient with the JaCoP plugin (72).

### Mass spectrometry analysis

#### Preparation of sucrose sedimented samples

Three independent biological replicates of untreated and arsenite-stressed cytosolic and non-cytosolic ultracentrifuged pellets were prepared from sequential detergent fractionation and sucrose sedimentation protocols described above, replacing iodoacetamide with chloroacetamide. Pellets were resuspended in triethylammonium bicarbonate (TEAB) buffer with 5% SDS (w/v), normalized (concentration determined by Bradford) to 10 μg of protein, and reduced using 10 mM DTT at 80C for 15 minutes using a thermomixer at 1,000 rpm. After cooling, samples were alkylated with 25 mM iodoacetamide in the dark for 30 min at room temperature. SDS was removed, and samples were digested with 1:15 mass ratio of Sequencing Grade Modified Trypsin (Promega) using an S-trap mini device (Protifi) according to manufacturer’s instruction (digestions at 47C for 1 hour). Recovered peptides were lyophilized to dryness and reconstituted at 0.5 μg/μL in 1% trifluoroacetic acid (TFA) and 2% acetonitrile (MeCN). Equal volumes of each sample were mixed to generate a study pool quality control (SPQC) sample.

#### Preparation of unenriched and K-GG enriched samples

Three independent biological replicates of untreated and arsenite-stressed cytosolic and non-cytosolic fractions were prepared from the sequential detergent fractionation protocol described above, replacing iodoacetamide with chloroacetamide. Samples were split into 5 subaliquots (200 μL each) and a chloroform-methanol precipitation was performed. A total of 4 mg of protein of each sample was aliquoted and normalized using 8M urea solution. Samples were reduced for 15 minutes at 80C and alkylated with 20 mM iodoacetamide for 30 minutes at room temperature. After further dilution with 2280 μL of 50 mM ammonium bicarbonate (AmBic), 60 μL of TPCK-treated trypsin (60 μg per sample) were added and samples were digested overnight at 32C and 1000 rpm. After the digestion, 30 μL of neat trifluoroacetic acid was added to a final concentration of 1% per volume. Sample clean-up was performed using Sep-Pak C18 cartridges (500 mg) according to the manufacturer’s instructions. Peptides were lyophilized overnight and an immunoaffinity purification was performed using the PTMScan® Ubiquitin Remnant Motif (K-ε-GG) Kit according to the manufacturer’s instructions with the following modifications: the 200 μL antibody aliquoted was diluted with 80 μL PBS, and 12 x 20 μL aliquots were prepared. Peptides were eluted in 2 x 50 μL of 0.15% TFA. SPQC samples were prepared from 2.5 μL of each sample (30 μL total). Four SPQC samples, and one 30 μL replicate of individual samples were loaded onto Evotip Pure tips.

#### Quantitative data-independent acquisition (DIA) LC-MS/MS analysis

Quantitiative LC/MS/MS of the sucrose sedimented samples was performed on 1.75 μL of each sample and three replicates of an SPQC pool, using a M-Class UPLC system (Waters) coupled to a coupled to a Fusion Lumos high resolution accurate mass tandem mass spectrometer (Thermo) via a Nanospray Flex Ion source. Samples were first trapped on a Symmetry C18 180 μm × 20 mm trapping column (5 μl/min at 99.9/0.1 v/v H2O/MeCN) followed by an analytical separation using a 1.7 μm ACQUITY HSS T3 C18 75 μm × 250 mm column (Waters) with a 90 min gradient of 5 to 30% MeCN with 0.1% formic acid at a flow rate of 400 nL/min and column temperature of 55°C. MS analysis used a staggered, overlapping window data-independent acquisition (DIA) method with a 120,000 resolution precursor ion (MS1) scan from 390-1010 m/z, AGC target of 1000% and maximum injection time (IT) of 60 ms and RF lens of 40%; data was collected in centroid mode. MS/MS was performed using tMS2 method with default charge state = 3, 15,000 resolution, precursor mass range of 400-1000, 151 x 8 m/z windows, AGC target of 1000%, maximum IT of 20 ms, a NCE of 30, with a 75-window cycle. An RF lens of 40% was used for MS1 and DIA scans.

Quantitative LC/MS/MS of the unenriched samples was performed on 3 μL of each sample diluted into 60 μL, along with replicates of an SPQC pool, using an Evosep One LC coupled to a Thermo Orbitrap Exploris 480 via a Nanospray Flex ionization source. The LC was interfaced to the MS using a PepSep sprayer and stainless steel (30 μm) emitter. The MS analysis used a 120,000 resolution precursor ion (MS1) scan from 480-920 m/z, AGC target of 1000%, and 60 ms maximum injection time (IT), collected every 0.6 s in centroid mode. MS/MS was performed using a DIA method with default charge state = 3, precursor mass range of 500-900 m/z, 16 m/z isolation windows, 30,000 resolution, AGC target of 1000%, maximum IT of 60 ms, a NCE of 30, and loop count of 25. An RF lens of 40% and centroid scans were used for MS1 and DIA scans.

Quantitative LC/MS/MS of the K-GG enriched samples was performed on ∼1/3 of each sample and three replicates of an SPQC pool, using an Evosep One LC coupled to a Thermo Orbitrap Astral via a Nanospray Flex ionization source. The LC was interfaced to the MS using a PepSep sprayer and stainless steel (30 μm) emitter. The MS analysis used a 240,000 resolution precursor ion (MS1) scan from 380-980 m/z, AGC target of 500%, and 50 ms maximum injection time (IT), collected every 0.6 s in centroid mode. MS/MS was performed using a DIA method with default charge state = 3, precursor mass range of 380-980, 10 m/z isolation windows, AGC target of 500%, maximum IT of 20 ms, and a NCE of 28. An RF lens of 40% was used for MS1 and DIA scans.

#### Quantitative analysis of all DIA data

Raw MS data was demultiplexed and converted to *.htrms format using HTRMS converter, and processed in Spectronaut 18 (Biognosys, 18.3.230830.50606). A spectral library was built using direct-DIA searches with a Homo sapiens database, downloaded from Uniprot and appended contaminant sequences using FragPipe. Search settings included N-terminal trypsin/P specificity up to 2 missed cleavages, and peptide length from 7-52 amino acids with the following modifications: carbamidomethyl (Cys), N-terminal protein acetylation, Met oxidation, and (for enriched samples only) GlyGly(K). For DIA analysis, default extraction, calibration, identification, and protein inference settings were used. Peptide and protein quantification were performed at MS2 level with q-value sparse settings (precursors that met a q-value<0.01 in at least one run were included for quantification). For visualization in Spectronaut, local normalization was performed and protein abundances were calculated using the MaxLFQ algorithm (73).

#### Statistical methods and data visualization

Statistical tests and visualizations were conducted using R version 4.3.0. Pairwise comparisons were assessed using the paired Student’s t-test, and unpaired comparisons were performed using the unpaired Student’s t-test. P-values < 0.05 were considered significant. Volcano plots and ordinal ranking plots were generated using the ggplot2 package. The specific enzyme lists were gathered from either the HUGO Gene Nomenclature Committee (E2s, DUBs), or UniProt codes gathered from their respective GO terms (E3s, ubiquitin binding proteins). The lists for ribosomal proteins were curated by matching the keywords “RPL” and “RPS” in the gene descriptions. The lists for subcellular proteins (cytosol, ER, mitochondria) were obtained from The Human Protein Atlas, and excluded proteins that were identified in multiple compartments. GO enrichment analysis was done using DAVID functional annotation clustering. Principal component analyses were performed in Spectronaut.

### Flow cytometry-based ROS assays

Cells with indicated treatments were incubated with CM-H2DCFDA (Thermo, C6827) for 15 minutes before harvesting by trypsinization. Cells were resuspended and PBS, and analyzed using a BD FACSCanto flow cytometer. Data was analyzed using FlowJo software. Cells were gated for live populations, then for singlets using forward and side scatter parameters. Then, median FITC fluorescence among live, singlet cell populations were used to determine ROS levels by sample.

### Puromycin incorporation assay

Cells were grown to 80% confluence and treated with the indicated stressors and/or inhibitors. Cells were then pulsed with puromycin (Gibco, A1113802) at a final concentration of 10 μg/mL 5 minutes before harvesting, then washed with PBS and trypsinized. Whole cell lysis protocol was followed as described above, and a western blot protocol was followed as described above probing for puromycin intensity as a proxy for translation activity at the time of harvest.

### Statistical analysis

Unless otherwise stated, all experiments displayed in this study were performed in biological triplicates. Analyses used Students t-test, and differences were considered significant with p value < 0.05. All statistical analyses (outside of proteomics data) were performed using GraphPad Prism 10.0.0.

## DATA AVAILABILITY

All LC-MS/MS proteomics data (.raw files) have been deposited to the MassIVE repository partnered with ProteomeXchange with the dataset identifier PXD052838. All other data are available in the associated supplemental data files. Further requests should be requested to the corresponding contact, Gustavo Silva.

## ACKNOWLEDGMENTS

We thank all members of the Silva lab, especially Shannon Dougherty, Géssica Barros, Clara dos Santos, Blanche Cizubu, Omur Kayikci, and Vanessa Simões, for technical assistance and constructive feedback. Additionally, we thank the Onishi and Magwene labs for sharing equipment and assistance. We also thank Chris Nicchitta and the Nicchitta lab for technical assistance, sharing equipment and resources, and constructive feedback. We also thank Lisa Cameron, Yasheng Gao, and the Duke Light Microscopy Core Facility for microscope support. We also thank Marlene Violette and the Duke Proteomics and Metabolomics Core Facility for assistance with data acquisition and analysis. We also thank the Duke Cancer Institute Flow Cytometry Core for flow cytometer support. Multiple figures in this manuscript were created using BioRender. This work was funded by National Institutes of Health grants R21ES032964 (G.M.S.), R35GM137954 (G.M.S.) and F30ES034271 (A.O.M.), Chan Zuckerberg Initiative SDL2022-253663 (G.M.S.), Burroughs Wellcome Fund Graduate Diversity Enrichment Program Award 1022058 (A.O.M.), ETH Fellowship 1-008383 (S.M.), and a Career Seed grant (S.M.).

## AUTHOR CONTRIBUTIONS

A.O.M. and G.M.S. conceived the project, designed experiments, interpreted results, and wrote the manuscript. A.O.M. performed all experiments. A.O.M., S.M., and M. W. F. analyzed data. All authors edited and contributed to its final version.

## ABBREVIATIONS

The abbreviations used are:

PTM: post-translational modification
ROS: reactive oxygen species
K63ub: K63-linked ubiquitin chains
K48ub: K48-linked ubiquitin chains
NaAsO_2_: sodium arsenite
VCP: valosin-containing protein
ER: endoplasmic reticulum
UBD: ubiquitin binding domain
RTU: redox control of translation by ubiquitin
RQC: ribosome-associated protein quality control
ERAD: ER-associated degradation
UPR: unfolded protein response
E2: ubiquitin conjugating enzyme
E3: ubiquitin ligase
DUB: deubiquitinase
ANS: anisomycin

