## Supplemental Figures S1-S6 for "Localized K63 ubiquitin signaling is regulated by VCP/p97 during oxidative stress"

### Supplemental Figure S1

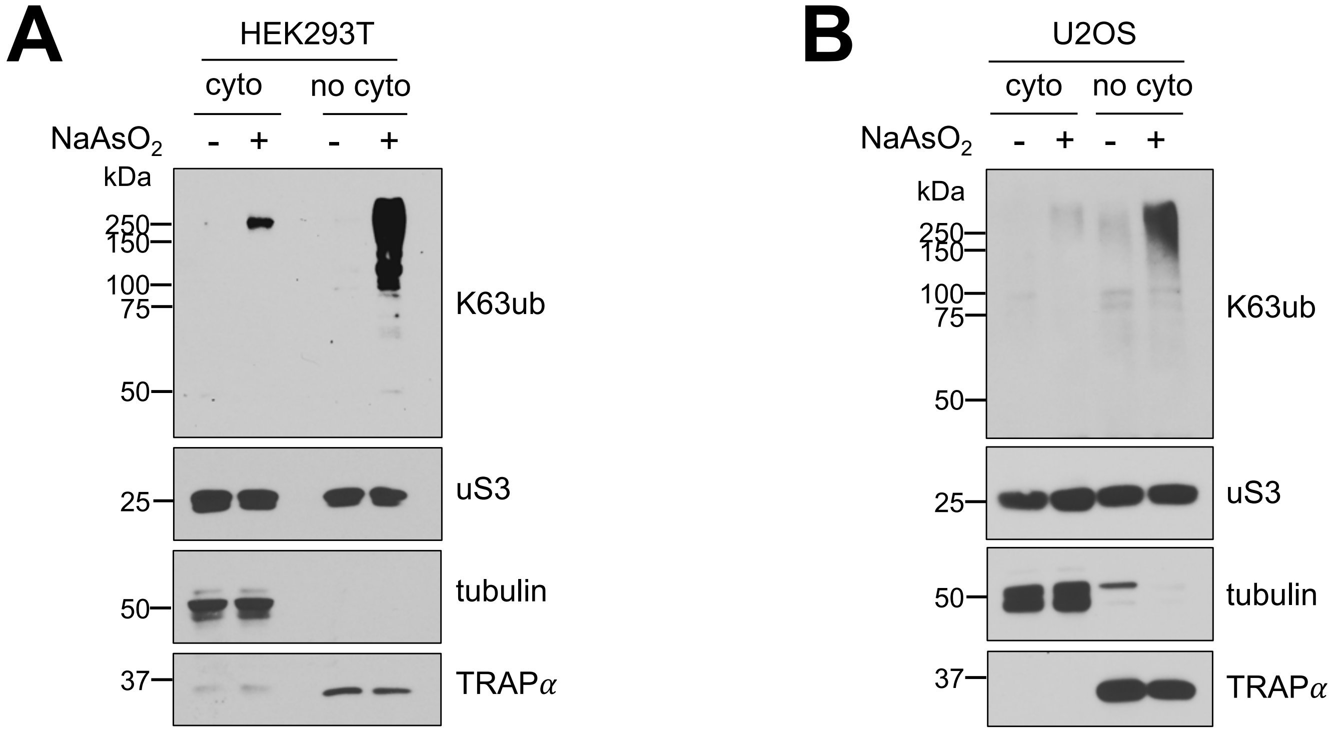


**Supplemental Figure S1. Arsenite stress induces non-cytosolic K63 ubiquitin accumulation in various human cell types.**

A and B – Immunoblots of anti-K63ub from purified fractions of either HEK293T or U2OS cells treated with 1 hour of 0.5 mM $NaAsO_2$. Anti-uS3 was used as a loading control for each purified fraction. Before ultracentrifugation, anti-tubulin and anti-TRAPα were used as fractionation controls for the cytosol and non-cytosolic fractions, respectively.

### Supplemental Figure S2

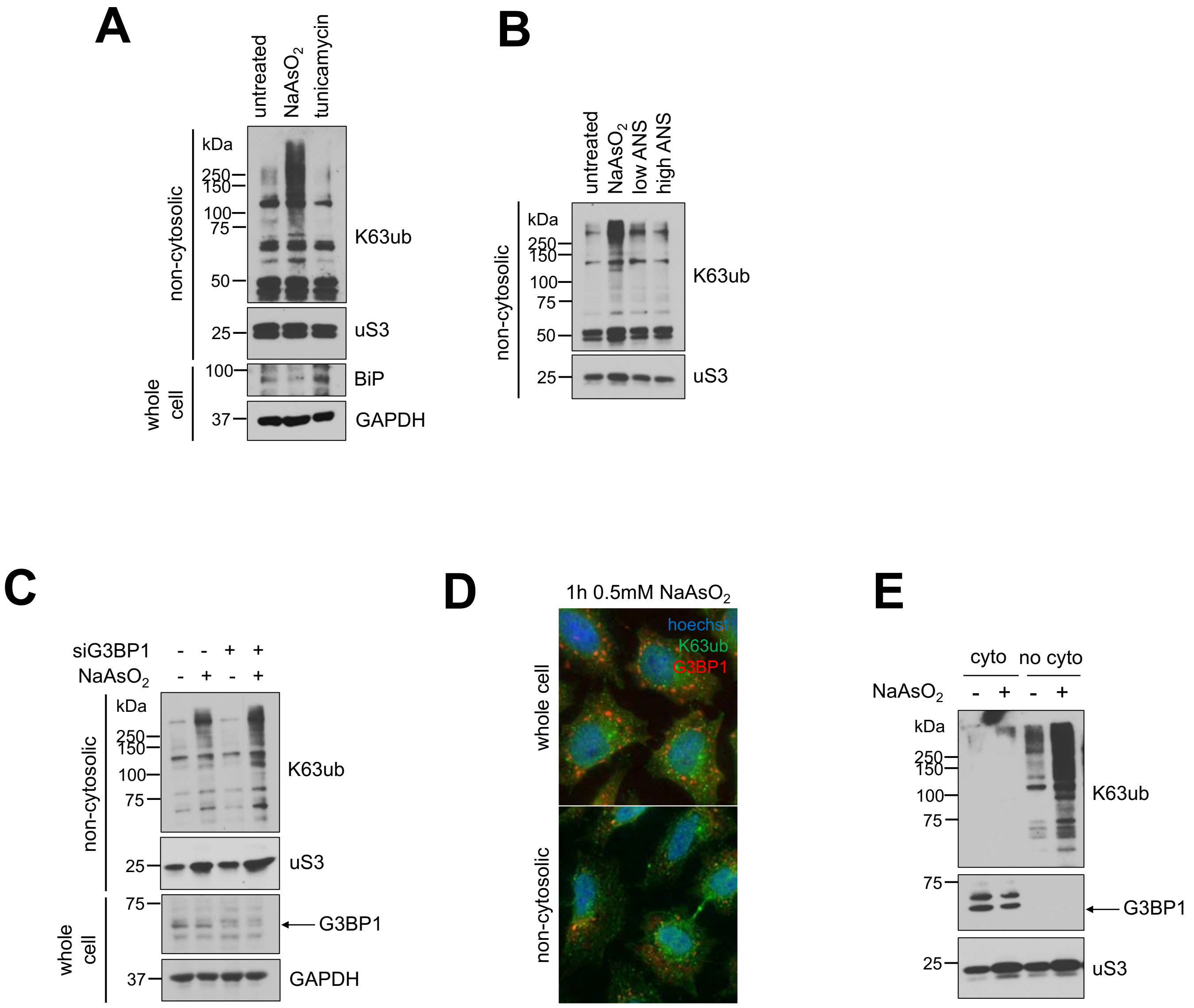


**Supplemental Figure S2. Other ubiquitin-mediated responses do not solely account for K63 ubiquitin accumulation.**

A-C – Immunoblots of anti-K63ub from non-cytosolic fractions of cells treated with 1 hour of 0.5 mM $NaAsO_2$, A) 15 minutes of tunicamycin (ER stress), B) low and high concentrations of anisomycin (ribosome collisions), or C) siG3BP1 (stress granule disassembly). Anti-uS3 was used as a loading control. In total cell lysates, successful induction of ER stress was measured through the unfolded protein response (UPR) and assessed using anti-BiP, and successful knockdown of G3BP1 protein levels were assessed with anti-G3BP1, both using anti-GAPDH as a loading control.

D – K63ub puncta present during arsenite stress does not colocalize with G3BP1. Immunofluorescence microscopy of anti-K63ub and anti-G3BP1 from cells treated with 1 hour of 0.5 mM $NaAsO_2$. Cells were prepared without and with a digitonin wash to release cytosolic components before fixation. Hoechst was used to identify cells. Scale bar = 10 $\mu$m.

E – Majority of G3BP1 under stress localizes to cytosolic compartments. Immunoblots of anti-K63ub and anti-K48ub from fractions of cells treated with 1 hour of 0.5 mM $NaAsO_2$. Anti-uS3 was used as a loading control for each purified fraction.

### Supplemental Figure S3

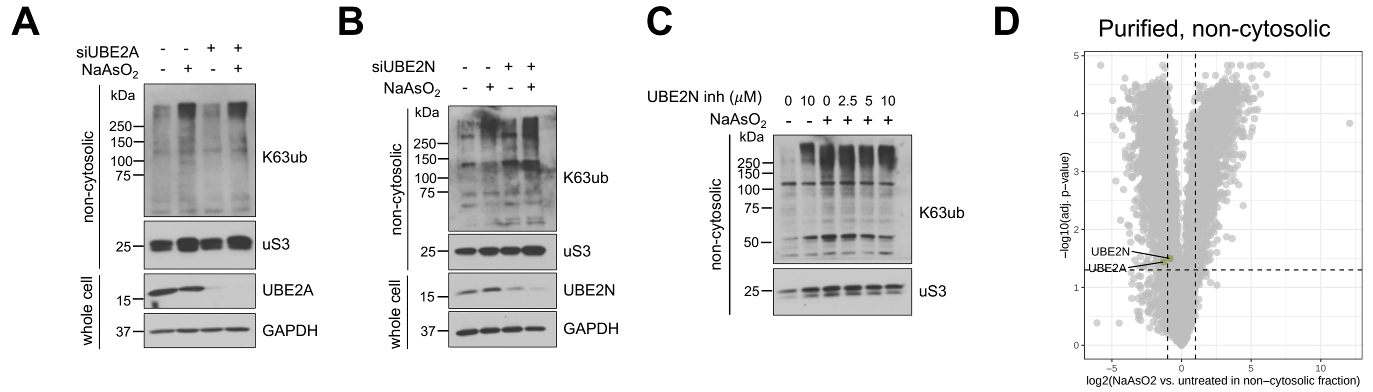


**Supplemental Figure S3. UBE2A and UBE2N do not solely account for K63 ubiquitin accumulation.**
A and B – Immunoblots of anti-K63ub from non-cytosolic fractions of cells transfected with siVCP versus siControl, then treated without and with 1 hour of 0.5 mM $NaAsO_2$. Anti-uS3 was used as a loading control. Successful knockdown of UBE2A or UBE2N protein levels were assessed from total cell lysates with anti-UBE2A and anti-UBE2N, using anti-GAPDH as a loading control.
C – Immunoblots of anti-K63ub from non-cytosolic fractions of cells treated with increasing concentrations of an inhibitor of UBE2N in the context of 1 hour of 0.5 mM $NaAsO_2$. Anti-uS3 was used as a loading control.
D – Volcano plots showing enrichments of proteins under arsenite stress in the non-cytosolic and cytosolic fractions, with UBE2A and UBE2N highlighted.

### Supplemental Figure S4

**A** Pre-enrichment

**B** Ubiquitinated peptides

**C** Purified compartments

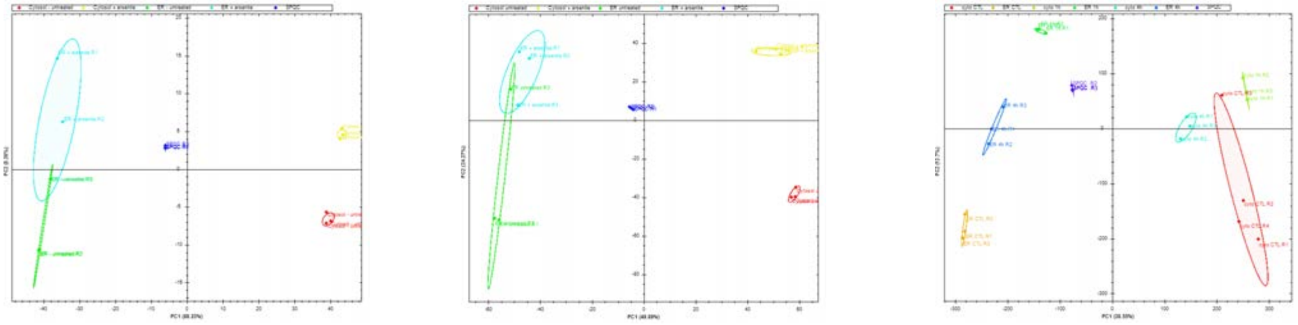

**Supplemental Figure S4. Variance between samples in proteomics datasets highlighted with principal component analyses.**
A-C – Principal component analyses from A) total abundance, B) ubiquitin peptide enriched, and C) sucrose sedimented datasets.

### Supplemental Figure S5

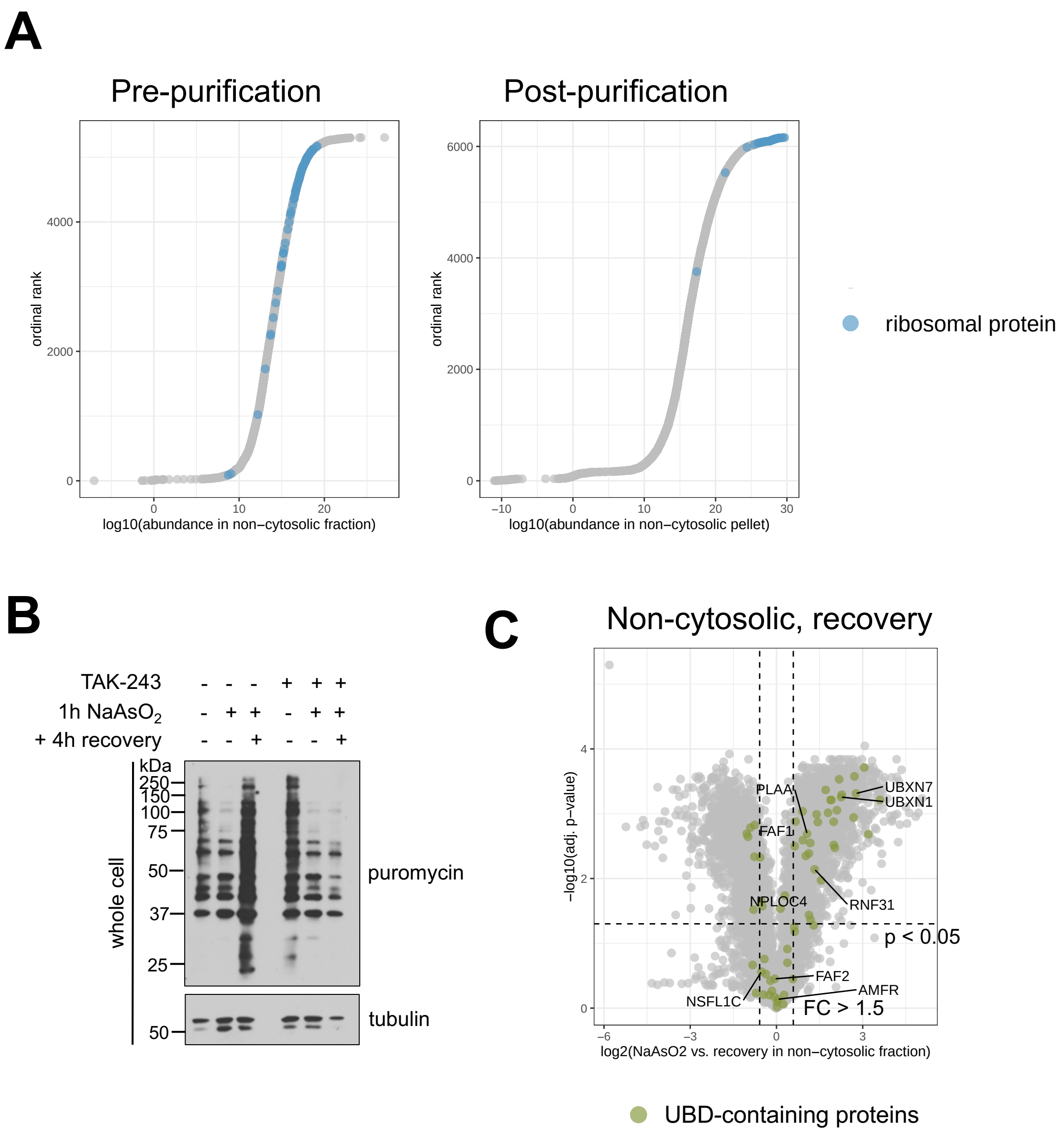


**Supplemental Figure S5. Proteomic analysis reveals additional details of purified non-cytosolic compartments under arsenite stress.**
A – Ordinal ranks of proteins in the non-cytosolic fraction before and after sucrose sedimentation. Highlighted proteins are large and small subunit ribosomal proteins.
B – Immunoblots of anti-puromycin from whole cell lysates of cells preincubated with or without 1 μM TAK-243, then 1 hour of 0.5 mM $NaAsO_2$ with or without a 4-hour washout. Anti-tubulin was used as a loading control.
C – Volcano plots showing enrichments under arsenite stress compared to recovery of proteins with ubiquitin binding domains in the non-cytosolic fractions. Known VCP adaptors were highlighted among the proteins with UBDs in both plots.

### Supplemental Figure S6

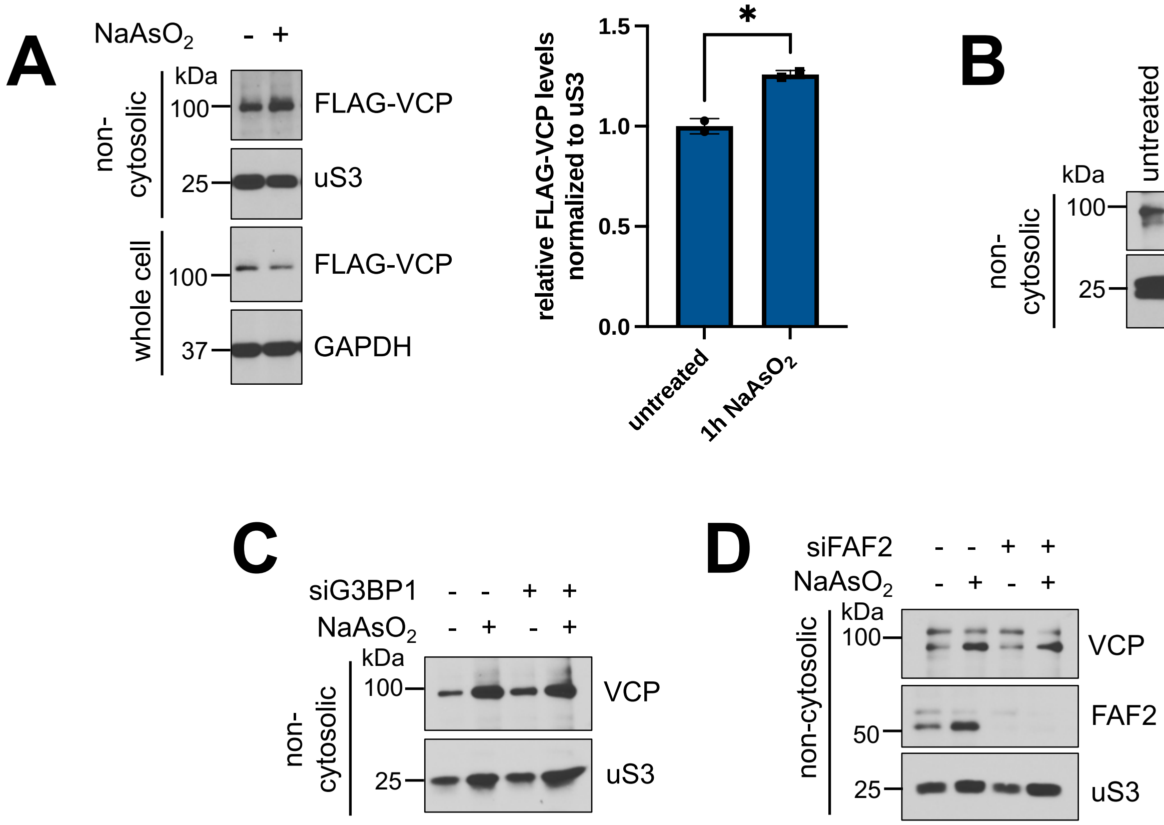


**Supplemental Figure S6. Other ubiquitin-mediated responses do not solely account for VCP accumulation.**

A – Arsenite stress induces exogenously expressed FLAG-VCP accumulation. Immunoblots of anti-FLAG from non-cytosolic fractions of cells transfected with FLAG-VCP plasmids for 24 hours, then treated the following day with 1 hour of 0.5 mM $NaAsO_2$. Anti-uS3 was used as a loading control for each purified fraction. Equal expression of FLAG-VCP were assessed under total cell lysate with anti-FLAG, using anti-GAPDH as a loading control. Densitometry analysis represents relative amount of FLAG-VCP normalized to uS3 loading control for two replicates.

B-D – Immunoblots of anti-VCP from non-cytosolic fractions of cells treated with 1 hour of 0.5 mM $NaAsO_2$, A) 15 minutes of tunicamycin (ERAD), B) siG3BP1 (stress granule disassembly), or C) siFAF2 (stress granule disassembly). Anti-uS3 was used as a loading control. In total cell lysates, successful induction of ER stress was measured through the UPR and assessed using anti-BiP, and successful knockdown of G3BP1 protein levels were assessed with anti-G3BP1, both using anti-GAPDH as a loading control. Anti-FAF2 was used to assess knockdown from the non-cytosolic fraction.
